# The natural selection of metabolism explains curvature in fossil body mass evolution

**DOI:** 10.1101/088997

**Authors:** Lars Witting

**Affiliations:** Greenland Institute of Natural Resources, Box 570, DK-3900 Nuuk, Greenland

**Keywords:** Evolution, metabolism, body mass, life history, allometry, fossil record

## Abstract

The natural selection of metabolism and mass can explain inter-specific allometries from prokaryotes to mammals (Witting 2017a), with exponents that depend on the selected metabolism and the spatial dimensionality (2D/3D) of intra-specific behaviour. The predicted 2D-exponent for total metabolism increases from 3/4 to 7/4 when the fraction of the inter-specific body mass variation that follows from primary variation in metabolism increases from zero to one.

A 7/4 exponent for mammals has not been reported from inter-specific comparisons, but I detect the full range of allometries for evolution in the fossil record. There are no fossil data for allometric correlations between metabolism and mass, but I estimate life history allometries from the allometry 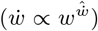 for the rate of evolution 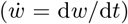 in mass (*w*) in physical time (*t*).

The 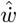 exponent describes the curvature of body mass evolution, with predicted values being: 3/2 (2D) for within niche evolution in small horses over 54 million years. 5/4 (2D) and 9/8 (3D) for across niche evolution of maximum mass in four mammalian clades. 3/4 (2D) for fast evolution in large horses, and maximum mass in trunked and terrestrial mammals. 1 for maximum mass across major life-forms during 3.5 billion years of evolution along a metabolic bound.

## 1 Introduction

Inter-specific allometries are essential for our understanding of natural selection. They reveal how the life histories of biological organisms evolve in correlations with mass, providing a fingerprint of the underlying natural selection cause.

One attempt to identify the cause explains metabolic scaling as a physiological adaptation where branching networks are optimised to supply the organism with energy for metabolism (West et al. 1997, 1999a,b; Banavar et al. 1999; Dodds et al. 2001; Dreyer and Puzio 2001; Rau 2002; Santillán 2003; Glazier 2010). This view is elaborated in the Metabolic theory of ecology, where physical and kinetic constraints on metabolism are influencing ecological processes like the rate of feeding and interaction (Gillooly et al. 2002; Brown et al. 2004; Sibly et al. 2012; Humphries and McCann 2014).

While physical and biochemical laws shape the ecology and physiology of biological organisms, it is the natural selection of metabolism and mass that causes the evolution of large organisms with metabolic scaling. Yet, this selection is not part of the allometric components of metabolic ecology, although the theory recognises the natural selection of mass in separate models on life history evolution (Brown and Sibly 2006; Bueno and López-Urrutia 2012).

These models use an adaptive fitness optimisation like a multitude of other studies that argue for a variety of intrinsic and ecological causes for the evolution of mass (e.g., McLaren 1966; Schoener 1969; Stanley 1973; Roff 1981, 1986; Stearns and Crandall 1981; Stearns and Koella 1986; Gould 1988; Maurer et al. 1992; Charlesworth 1994; Bonner 2006; Clauset and Erwin 2008; Smith et al. 2010; Charnov 2011; DeLong 2012; Shoemaker and Clauset 2014; Baker et al. 2015). Basically all these hypotheses assume constant relative fitnesses, which implies a frequency-independent selection with an increase in the rate of population dynamic growth (*r*) and/or carrying capacity (*k*). Fisher (1930) used this increase to formulate his fundamental theorem of natural selection (Witting 2000), a theorem that later became the cornerstone of *r/k* selection theory (Anderson 1971; Charlesworth 1971; Roughgarden 1971; Clarke 1972). But body masses that are selected by an increase in *r* and/or *k* does not explain the observed allometries, where an increase in mass is associated with a decline in *r* and *k* (Fenchel 1974; Damuth 1981, 1987).

A frequency-dependent selection of mass, on the other hand, is able to reconcile the observed inter-specific decline with an increase in mass (Simpson 1953; Dawkins and Krebs 1979; Parker 1979, 1983; Haigh and Rose 1980; Maynard Smith and Brown 1986; Vermeij 1987; Witting 2000). This brings us to the theory of Malthusian relativity (Witting 1995, 1997, 2008, 2017a,b), where the density-frequency-dependent selection of mass and metabolism explains the body mass exponents of allometries for fitness related traits like net assimilated energy (*ϵ*), mass-specific metabolism (*β*), life-periods (*t*), reproduction (*R*), survival (*p*), population growth (*r*), abundance (*n*), and home range (*h*).

### 1.1 The fingerprint of metabolic selection

Malthusian relativity defines body mass by the net assimilated energy that is allocated into size instead of being used in metabolism or allocated to offspring. This mass is measured by the energy that is released from combustion, with an approximate proportional conversion between mass in joules and mass in grams. The theory finds that the density-frequency-dependence of interactive foraging is generating an intra-specific population dynamic feedback that selects for non-negligible body masses, as illustrated in Figure 1. The feedback selects net assimilated energy into mass by a resource bias from interactive competition, with the ecological constraints—on the population-wide geometrical packing of foraging in home ranges and territories—selecting for an allometric scaling between mass and other fitness related traits (Witting 1995, 2017a).

**Figure 1:**
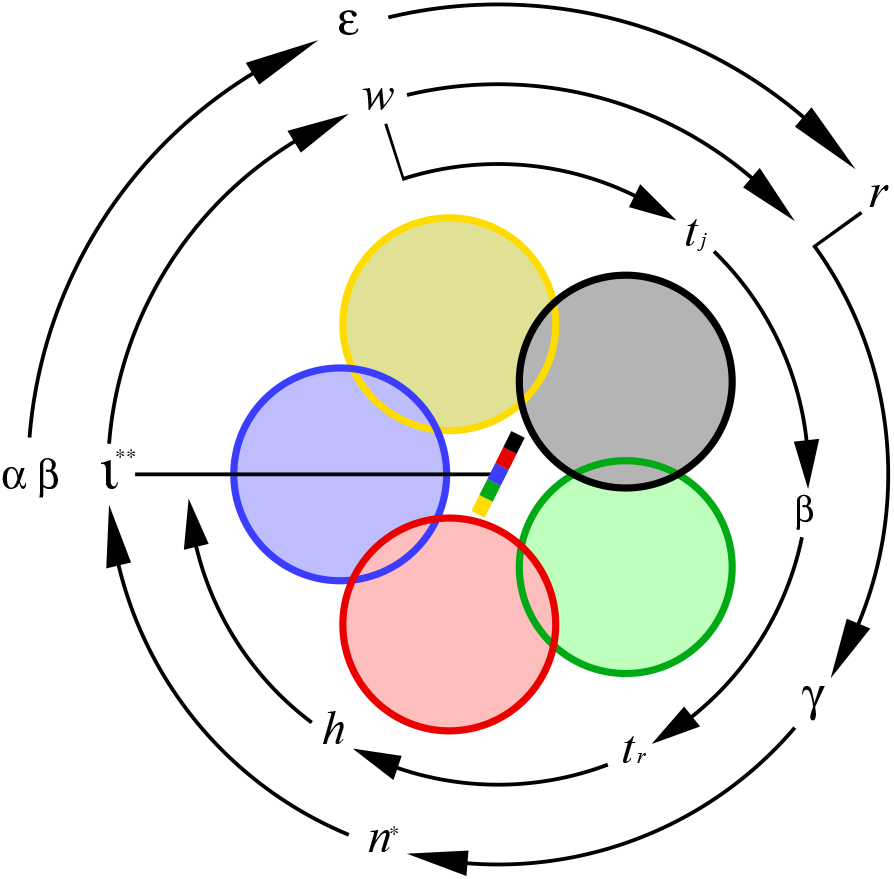
Population dynamic feedback selection. A diagram of the population dynamic feedback selection of Malthusian relativity, with symbols that relate to the population average, and coloured circles that symbolize individual home ranges in two-dimensional space with interactive competition in zones of overlap. The winners (dominating colour) of interactive competition monopolize resources, and this generates a body mass biased resource access that is proportional to the slope of the multi-coloured bar in centrum, with the invariant interference (*ι*) of the selection attractor (**) determining the selection of this bias. The primary selection on resource handling (*α*) and mass-specific metabolism (*β*) generates an exponential increase in the net assimilated energy (*ϵ*), and this maintains relatively high population dynamic growth (*r*) and a continued feedback selection for an exponential increase in mass (*w*). The feedback attractor is illustrated by the outer ring of symbols [*r*:population growth → *γ*:density regulation → *n*\*:population abundance → *ι*:interference level → *w*:selection on body mass → *r*:population growth]. Selection for a change in mass initiates the inner loop of mass-rescaling selection [*w*:mass → *t_j_*:juvenile period → *β*:metabolic rate → *t_r_*:reproductive period → *h*:home range → *ι*:interference]. Modified from Witting (2017b).

This mass-rescaling selection is caused by the natural selection of mass (Witting 2017a), and it is illustrated in Figure 1 by the inclusion of the inner loop of symbols. The result is a set **x** = {*ϵ, β,t, R,p,r,n, h*} of traits that evolve partial allometric correlations 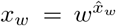 with mass (*w*), where *x_w_* ∈ **x** with allometric exponents 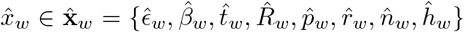, with subscript *w* denoting mass-rescaling selection. When the population ecological constraints are formulated in mathematical equations, we can express the different traits by their allometric functions of mass, and solve the equation system for the unknown values of the exponents 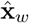. This solution (Witting 1995) includes the well-known 1/4 and 3/4 exponents of Kleiber (1932) scaling from a two-dimensional (2D) packing of the foraging pattern, with alternative 1/6 and 5/6 exponents being selected by a three-dimensional (3D) packing.

The natural selection of allometries, however, is more complex because the primary selection of metabolism generates net energy for the natural selection of mass (Witting 2017a,b). This is because the net assimilated energy (*ϵ*, SI unit J/s) is a product 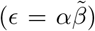 between the mechanical/biochemical handling of resource assimilation (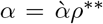; resource handling in short; SI unit J; 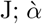: intrinsic handling; *ρ*\*\*:resource density at population equilibrium), and the pace of handling (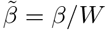; SI unit 1/s), where pace is selected as a proxy for the rate of mass-specific metabolism (*β*; SI unit J/Js), with *W* (SI unit J/J) defined as the mass-specific work of handling from one joule metabolised per unit body mass (Witting 2017a).

The primary selection of metabolism imposes a metabolic-rescaling on the rate dependent traits (Witting 2017a). This rescaling is then transformed into partial correlations of metabolic-rescaling allometries (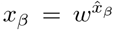, subscript *β* denotes metabolic-rescaling), i.e., the traits correlations with mass that evolve from metabolic-rescaling and the mass that is selected from the net energy that follows from the primary selection of mass-specific metabolism. The final allometries 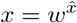 are products *x* = *x_β_x_w_* of the two partials of mass and metabolic rescaling selection. This implies exponents 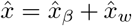 that depend on the relative importance of mass-specific metabolism for the evolution of net energy and mass, as described by a metabolic-rescaling exponent for mass-specific metabolism 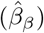 that varies from zero to one (Witting 2017a); with an associated final 2D exponent for mass-specific metabolism that varies from −1/4 to 3/4.

Instead of focussing on the exponent of the final allometry, i.e., 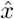 in 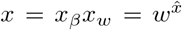, we may interpret the metabolic-rescaling component 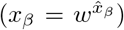 as the intercept of the mass-rescaling allometry 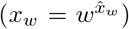. A well-known case of this interpretation is the difference in the intercepts of the allometries for ectotherm and endotherm vertebrates. Both taxa have Kleiber-like mass-rescaling exponents across the species within each taxon, yet the allometric intercept for mass-specific metabolism is higher in endotherms. The higher intercept implies a higher mass-specific metabolism for similar sized species, reflecting that metabolism is most essential for resource consumption in endotherms.

For vertebrates with a similar mass-specific metabolism, endotherms have a larger downscaling of metabolism by a stronger mass-rescaling component, i.e., a larger mass as the mass-rescaling exponents are about the same. This illustrates that endotherms tend to have larger body masses than ectotherms because they generate more net energy from a higher rate of resource handling.

The incorporation of primary selection on metabolism can explain also a wider range of allometries across lifeforms from virus over prokaryotes and larger unicells to multicellular animals. The exponents of empirical allometries have been found to change in transitions across the major taxa (e.g., Makarieva et al. 2005, 2008; DeLong et al. 2010), and Witting (2017a,b) was able to shows how a directional change—in the selection of metabolism and mass with an increase in mass—is explaining the evolution of the different allometries and lifeforms.

### 1.2 Allometries in time

Inter-specific allometries are usually used to describe relationships across existing species, and the study of Witting (2017a,b) is no exception. With this paper I extend the analysis to describe the natural selection of metabolism, mass, and allometries in time. This reveals potential natural selection causes behind some of the best-documented evolutionary trajectories in the fossil record. They describe body mass evolution at different scales, ranging from within-lineage evolution over 54 million years in horses (MacFadden 1986; Shoemaker and Clauset 2014), over maximum mass evolution in several clades of mammals (Smith et al. 2010; Okie et al. 2013), to the evolution of maximum mass across major lifeforms over 3.5 billion years (Bonner 1965; Payne et al. 2009).

In across species comparisons it is the amount of inter-specific variation in resource handling relative to the amount of variation in the primary selected component of mass-specific metabolism that determines the values of the selected allometric exponents. Yet, when we study the allometries of life history evolution in time, it follows that the allometric exponents that are selected at any given time will depend on the relative rates of evolution in the different traits. The selected rates of increase in net energy (*r_ϵ_* = dln *ϵ/dτ; τ*:per-generation time) and mass (*r_w_* = dln*w*/d*τ*) will depend on the selected rates of increase in resource handling (*r_α_*) and mass-specific metabolism (*r_β_β__*, with sub-subscript *β* denoting the primary selected component of metabolism). This implies that the selected allometric exponents dependent on the relative importance of metabolism for the selection of mass, as expressed e.g. by the *r_β_β_/r_α__*-ratio.

We may thus use the allometric exponents of an evolutionary body mass trajectory in time to determine the *r_β_β__/r_α_*-ratio of its underlying natural selection cause. However, to apply this method to fossils we need to solve the problem that the only life history trait that is commonly estimated for fossil animals is mass. How can we estimate the exponents of the underlying allometries, when there are no estimated traits to correlate with the observed evolutionary changes in mass?

I show that we can use the curvature in the rate of evolutionary change in mass to estimate the allometries and the underlying importance of mass-specific metabolism for the natural selection of mass. The rate of change in the mass of an evolving lineage can be expressed as an allometric function of mass

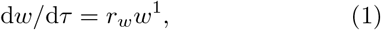

with the predicted exponential increase on the per-generation time-scale of natural selection (Witting 1997, 2003, 2018) being a log-linear increase with an allometric exponent of unity.

When the allometry for the rate of change in mass is determined empirically from fossils it is expressed in physical time, where

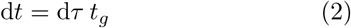

and

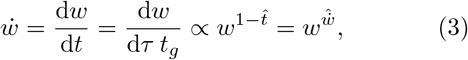

with *τ* being time in generations, *t* time in years, 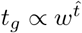 one generation in years at generation *τ*, 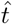 the allometric exponent for the evolution of generation time with mass within the evolutionary lineage over time, and 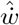 an allometric bending exponent that reflects the curvature of the rate of change in mass in physical time.

The log-linear trajectory of eqn 1 bends into a curve in physical time 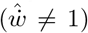 whenever 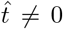, i.e., whenever the time-scale of natural selection evolves with the evolutionary changes in mass (Witting 1997; Okie et al. 2013). When natural selection time dilates by a generation time that increases with mass 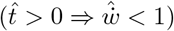 it bends body mass evolution downwards in a concave trajectory with a continuously declining dln *w*/d*t* derivative. Upward bending, with a convex trajectory and a continuously increasing derivative, occurs when the time-scale of natural selection contracts from a generation time that declines with mass 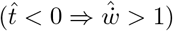.

I show theoretically that this bending—of a log-linear body mass trajectory on the time-scale of natural selection into a curved trajectory in physical time—is a fingerprint of the underlying primary selection of mass-specific metabolism. With an overall allometric scaling that depends on metabolic-rescaling, I find the curvature of body mass evolution to depend on the primary selected mass-specific metabolism, with the equations of natural selection predicting the 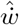 exponent of eqn 3 as a continuous function of the *r_β_β__/r_α_*-ratio.

In applying the theory to fossils I do not simply fit the model to trajectories, as this would just estimate the *r_β_β__/r_α_*-ratio with no direct first-principle predictions to be tested by observation. I focus instead on four simple and distinct types of natural selection that can be identified by first-principle arguments to specific trajectories in the fossil record, with each type of selection having its own theoretical 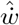 exponent and *r_β_β__/r_α_*-ratio. The intriguing question is then whether the observed exponents coincide with the first-principle prediction.

## 2 Methods

### 2.1 Theory

I extend the selection equations of Malthusian relativity to predict the bending exponent w for body mass evolution. My starting point is the evolutionary steady state with selection for an exponential increase in net assimilated energy and mass. The original version of the steady state (Witting 1997) did not partition the selection of net energy into an independent selection of the handling of resource assimilation and an independent selection of the pace of this process, as represented by mass-specific metabolism. But, with allometries that depend in part on the primary selected metabolism (Witting 2017a), a distinction is necessary to predict the bending exponents of evolutionary trajectories.

Appendix A develops the exponential increase of the evolutionary steady state with primary selection on handling and mass-specific metabolism (model symbols in Table 2). The resulting equations are combined with the results from Witting (2017a) and solved in Appendix B to obtain the allometric exponents as a function of the *r_β_β__/r_α_*-ratio. The theoretical exponent for generation time 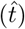 is then inserted into eqn 3 to calculate the bending exponent 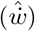 for body mass evolution in physical time.

The first of the four selection types that I consider for data confirmation is an unconstrained symmetrical selection across ecological niches. It generates a *r_β_β__/r_α_*-ratio of unity, with similar rates of increase in resource handling and mass-specific metabolism. Selection across niches may also cause a second non-symmetrical type of selection that is characterised by a *r_β_β__/r_α_*-ratio that approaches zero. The latter may occur when fast improvements in resource handling are easy, and the maximum size is increasing fast by an increase in resource handling that outruns the background evolution of mass-specific metabolism. The underlying cause may be a resource density that increases across niches, or a resource handling efficiency that increases as a mechanistic function of the evolutionary increase in mass.

These across niche selections are likely to occur in the larger species of clades that diversifies in evolutionary radiations. Not only are the larger species likely to evolve from adaptations to ecological niches that allow for a larger resource consumption, but nor should their niche access be limited by competitively superior species. This contrasts to smaller species that may have their niche access and resource handling constrained by the inter-specific dominance of larger species. The trajectory for the maximum size of a clade is the obvious place to look for an unconstrained evolution of mass.

The third selection relates to the alternative scenario where a lineage evolves within a relatively stable ecological niche. This results in an adaptation that selects resource handling to an evolutionary optimum where *r_α_* → 0 and the *r_β_β__/r_α_*-ratio approached infinity.

The final selection that I will examine relates to macroevolution on the time-scale of major transitions; first from prokaryotes to unicellular eukaryotes, then to multicellular animals, and then between the major lifeforms of multicellular animals that have taken the record of maximum size over time. Each of these transitions require a major reorganisation of the phenotype, and this implies a major restructuring of the biochemical/mechanical/ecological mechanisms of resource handling (*α*). The rate of evolution in this macroevolutionary component of *α* should thus be much smaller than the rates that are realised when a given lifeform adapts, by relatively simple phenotypic adjustments, to resources across ecological niches.

We may thus expect that the rate of macroevolutionary increase in resource handling will be much smaller than the persistent primary selected increase in mass-specific metabolism (*r_α_* ≪ *r_β_β__/r_α_*), so that the primary selected increase in metabolism should be able to outrun the decline from mass-rescaling on the macro evolutionary time-scale. Macroevolution should thus take mass-specific metabolism to an upper limit, as measured in the species with the highest metabolism, i.e., creating an invariant mass-specific metabolism across the smallest species of each lifeform. The result is macroevolution along a metabolic bound where the primary selected increase in metabolism is exactly outbalancing the mass-rescaling decline, i.e., where the product of metabolic-rescaling and mass-rescaling is zero 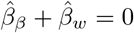.

Each of these four types of selection have a specific *r_β_β__/r_α_*-ratio; this ratio, however, is generally not constant on a microevolutionary time-scale. The realised rates of increase in *α* and *β* will depend on available mutations, and these are generally not homogeneous in time. But when integrating over longer time-scales we expect the *r_β_β__/r_α_*-ratio to stabilise at a central value. The four modes of selection can therefore not predict specific rates of increase at discrete moments in time, but only an integrated average rate over a longer time span.

The variation in the rates of evolution will also depend on fluctuations in climate and inter-specific competition, and the overall selection for size increase may shift to a decline if the net assimilated energy is declining due to an environmental or inter-specific competitive crisis (Witting 1997, 2008, 2018). Hence, we expect that the general trend for an increase in size is somewhat hidden in the fossil record; hidden behind a diverse and complex distribution of step-wise increasing and declining body masses.

### 2.2 Data

The expectation of a complex size distribution on top of a trend for a general increase agrees with mammalian data for the past 100 million years of evolution. They show a widespread increase in size (Alroy 1998; Baker et al. 2015) in agreement with Cope’s (1887) rule. Around 70% of descendant species are larger than their ancestors (Baker et al. 2015), about 30% are smaller, and the distribution of long-term trajectories is far from a homogeneous exponential-like increase in mass.

This paper does not apply the evolutionary steady state to the diversity of trajectories in the fossil record, and nor does it attempt to estimate selection on microevolutionary time-scales. It is only trying to characterise the average long-term forms of selection for increased size for cases where the fossil record show a relatively homogeneous increase in size over time spans that are so long that the general increase is evident in relation to the variation of the data. I am thus focusing on a few long-term trajectories that all have a relatively smooth exponential-like increase in size, as predicted by the evolutionary steady state.

In agreement with expectations, the maximum size of a clade is often found to increase in a near-exponential manner during periods of phylogenetic radiation (e.g., Hayami 1978; Trammer 2005; Smith et al. 2010; Okie et al. 2013). Hence, I use the data from Smith et al. (2010) and Okie et al. (2013) on the evolution of maximum mass over millions of years in mammals and mammalian clades to resemble evolution across ecological niches.

It is essential to note that these trajectories of maximum mass are inherently connected with speciation during evolutionary radiation, and they do generally not represent the continued evolution of a single lineage. They represent instead the overall time-bounds on the evolution of potential resource handling given an assumed relatively constant rate of evolution in mass-specific metabolism across the larger species in the clade. The macroevolutionary estimates of the bending exponent are thus integrating across species to reflect the overall trend in the evolutionary variation in mass-specific metabolism and maximum mass within a clade.

Another case is evolution in fossil horses (Equidae) over the past 57 million years in North America, which is one of the best documented cases for an exponential-like increase in body mass (MacFadden 1986; Witting 1997; Shoemaker and Clauset 2014). During the Eocene and Oligocene, from about 57 to 25 million years ago (ma), horses were browsers with a relatively small increase in size (from about 25 to 50 kg) and an absence of a strong diversification of species. In the early and middle Miocene from about 25 to 10 ma horses had a major pulse of body mass evolution, with an associated strong taxonomic diversification of both browsing and grazing horses (MacFadden 1986). This lead to the evolution of large horses (up to about 1 ton for *Equus giganteus* around 0.012 ma, Eisenmann 2003), yet the evolution of small types continued with the 80 kg *Nannippus peninsulatus* around 3.5 ma being an example of the latter.

In order to capture this pattern, I divide the MacFadden (1986) data on body masses of fossil horses into small (below 100 kg) and large horses (above 100 kg) to consider their evolution separately. Small browsing horses are likely to represent well-adapted lineages that evolve within a restricted niche space where *r_α_* is about zero and the *r_β_β__/r_α_*-ratio approaches infinity. Yet, we expect a transition where the diversification of larger horses after 25 ma reflects selection across niches. This would imply a *r_β_β__/r_α_*-ratio around unity, if not zero as the fast evolution of size indicates a resource handling that might outrun the background selection of mass-specific metabolism.

Finally, I use data from Bonner (1965) and Payne et al. (2009) on the evolution of maximum size in mobile organisms over 3.5 billion years, to test for a macroevolutionary invariance in mass-specific metabolism across major lifeforms. The result of this analysis is doublechecked against the potential existence of an invariance in the mass-specific metabolism of extant species across the major lifeforms of prokaryotes, unicellular eukaryotes, and the multicellular animals of aquatic ectotherm, terrestrial ectotherms, and terrestrial endotherms. This macroevolutionary invariance is already documented for species distributions as a whole (Makarieva et al. 2005, 2008; Kiørboe and Hirst 2014; Witting 2017b), and I use the data of Savage et al. (2004) and Makarieva et al. (2008) to select the three species with the highest mass-specific metabolism from each group to test for the existence of a mass invariance at the upper metabolic limit (Supplementary Table S7).

Having estimates of mass over time for each trajectory, I calculate rates of evolution d*w*/d*t* = (*w_t_d__* − *w_t_a__*)/(*t_d_* − *t_a_*) from adjacent data points at time *t_a_* and *t_d_* (Supplementary Tables S1 to S6; *w_t_a__* is the ancestral, and *w_t_d__* the descendant, mass. For horses, I use the ancestral-descendant species pairs that were identified by MacFadden 1986). The bending exponent 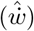 is then estimated as the slope of the linear regression of ln(d*w*/d*t*) on ln 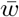 for cases with five or more data points 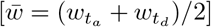.

Focussing on the natural selection of increased size, I include only species comparisons where descendants are larger than ancestors in my estimation of bending exponents. This will give me the best estimates of the underlying process, as it removes cases where a climatic and/or inter-specific competitive fluctuation is causing a short-term decline in the size of a taxa that is otherwise increasing steadily over time.

The predicted bending exponents will not only depend on the *r_β_β__/r_α_*-ratio, but also on the dimensionality of the interactive foraging behaviour. This should reflect the dominant spatial packing of the home ranges and territories that the individuals of a species compete for. For this classification I follow the data based system in Witting (2017a), where the dimensionality of different taxa follows from the average point estimate of the allometric exponent for mass-specific metabolism. These estimates agree in most cases with first-principle expectations, where the individuals of many pelagic and tree living species (like cetaceans and primates) classify as 3D, with an extra vertical dimension relative to most mammals that compete for 2D home ranges only.

## 3 Results

### 3.1 Exponential increase

Appendix A finds that primary selection on resource handling (*α*) and mass-specific metabolism (*β*) generates an exponential increase on the per-generation time-scale in the average resource handling, mass-specific metabolism, and net assimilated energy (*ϵ*) of the individuals in the population. The rate of exponential increase in net energy

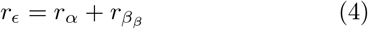

is the sum of the rates of increase in handling (*r_α_*) and mass-specific metabolism (*r_β_β__*).

The exponential increase in net energy generates a sustained population dynamic feedback selection. The net energy that is allocated into reproduction causes population growth and interactive competition, with the latter selecting net energy into mass

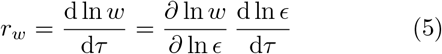

at the cost of a continued increase in the reproduction and density of, and interactive competition within, the population. The result is an evolutionary steady state, where an allometric selection *∂* ln *w/∂* ln 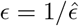 selects an exponential increase in mass

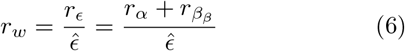

from the allometric exponent for net energy 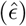 and the underlying rates of increase in resource handling and mass-specific metabolism.

This selection of mass is associated with a mass-rescaling selection (Witting 2017a) that dilates the time-scale of reproduction (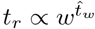, with 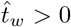) by a partial decline in mass-specific metabolism 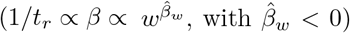. This dilation of natural selection time maintains a balance between net energy and mass that maintains the population dynamic pressure on the interactive selection of net energy into mass. A larger offspring metabolises more energy during growth, and this will cause a decline in average reproduction (and population growth and interactive competition) with an increase in mass, unless the rate of mass-specific metabolism is selected to decline and the reproductive period is selected to increase (Witting 2017a). The result is mass-rescaling allometries, where exponents like

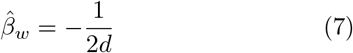

were shown by Witting (1995) to follow from the spatial dimensionality (*d*) of the interactive foraging behaviour.

### 3.2 Allometries in time

The partial decline in mass-specific metabolism from mass-rescaling selection occurs independently of the primary selection for an exponential increase in mass-specific metabolism. The latter generates a metabolic-rescaling of the life history, with Appendix B showing that the evolutionary steady state implies a metabolic-rescaling exponent for mass-specific metabolism

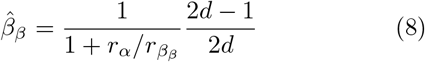

that depends on the *r_β_β__/r_α_*-ratio and ecological dimensionality (*d*). Combined with the mass-rescaling exponent of eqn 7 it defines the final allometry 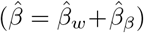 as illustrated in Fig. 2.

**Figure 2:**
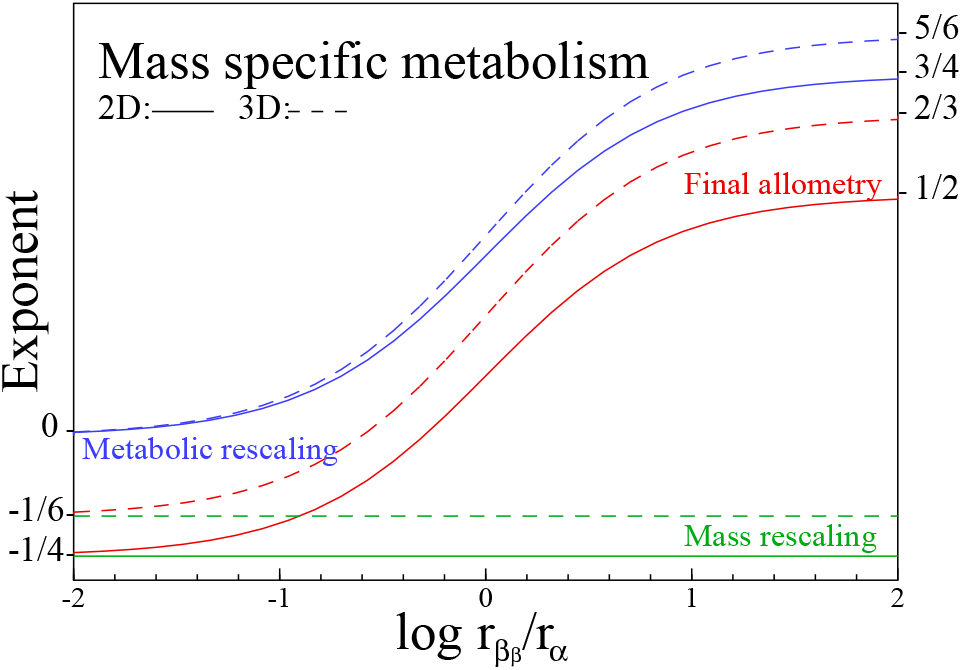
**The allometric scaling** of mass-specific metabolism 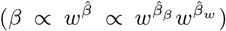 as it evolves from the *r_β_β__/r_α_*-ratio given 2D and 3D interactions. The blue curves are the exponents of metabolic-rescaling 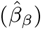, the green lines the exponents of mass-rescaling 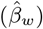, and the red curves the exponents of the final allometries 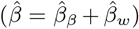.

A constant mass-rescaling intercept 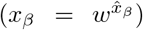 with no metabolic-rescaling 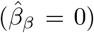 is obtained only when there is no primary evolution in mass-specific metabolism and *r_β_β__/r_α_* = 0. More generally, the mass-rescaling intercept will increase with an evolutionary increasing body mass, because the mass increase follows partly, or fully, from an evolutionary increase in mass-specific metabolism. For symmetrical evolution where *r_β_β__/r_α_* = 1, the mass-rescaling intercept will increase to the 3/8 power of body mass in 2D, and the 5/12 power in 3D. At the limit where metabolism evolves unconstrained (*r_β_β__* > 0) with a constraint on resource handling (*r_α_* = 0, with *r_β_β__/r_α_* ≈ to), we find a mass-rescaling intercept for mass-specific metabolism that scales to the 3/4 power in 2D, and the 5/6 power in 3D.

Having the metabolic-rescaling exponents for mass-specific metabolism as a function of the *r_β_β__/r_α_*-ratio, it is straightforward to transform the theoretical exponents from Table 3 in Witting (2017a) into the allometric exponents of Table 1. For the body mass trajectory of an evolutionary lineage in time, it gives the allometric exponents of the different traits as a function of the *r_β_β__/r_α_*-ratio and the dimensionality of the foraging behaviour.

**Table 1:** Theoretical allometries. Allometric exponents for exponential body mass evolution on a per-generation time-scale, as they evolve from allometric rescaling with selection on mass and mass-specific metabolism. The exponents depend on the dimensionality of the interactive behaviour, and on the ratio of the exponential rate of increase in mass-specific metabolism and resource handling (*r_β_β__/r_α_*). **Symbols:** *ϵ*:net energy; *α*:resource handling; *β*:mass-specific metabolism; *t*:biotic periods in physical time; *p*:survival; *R*:lifetime reproduction; *r*:population growth rate; *h*:home range; *n*:population density; 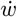:rate of body mass evolution in physical time (accent hat denotes exponent).

| a) One-dimensional interactions ($\hat{\epsilon} = 1/2$) | | | | | | | | | |
| --- | --- | --- | --- | --- | --- | --- | --- | --- | --- |
| $\frac{r_{\beta\beta}}{r_{\alpha}}$ | $\hat{\alpha}$ | $\hat{\beta}$ | $\hat{t}$ | $\hat{p}$ | $\hat{R}$ | $\hat{r}$ | $\hat{h}$ | $\hat{n}$ | $\hat{w}$ |
| 0 | $\frac{1}{2}$ | $-\frac{1}{2}$ | $\frac{1}{2}$ | 0 | 0 | $-\frac{1}{2}$ | 1 | $-\frac{1}{2}$ | $\frac{1}{2}$ |
| 1 | $\frac{1}{4}$ | $-\frac{1}{4}$ | $\frac{1}{4}$ | $\frac{1}{4}$ | $-\frac{1}{4}$ | $-\frac{1}{4}$ | 1 | $-\frac{3}{4}$ | $\frac{3}{4}$ |
| ∞ | 0 | 0 | 0 | $\frac{1}{2}$ | $-\frac{1}{2}$ | 0 | 1 | -1 | 1 |

| b) Two-dimensional interactions ($\hat{\epsilon} = 3/4$) | | | | | | | | | |
| --- | --- | --- | --- | --- | --- | --- | --- | --- | --- |
| $\frac{r_{\beta\beta}}{r_{\alpha}}$ | $\hat{\alpha}$ | $\hat{\beta}$ | $\hat{t}$ | $\hat{p}$ | $\hat{R}$ | $\hat{r}$ | $\hat{h}$ | $\hat{n}$ | $\hat{w}$ |
| 0 | $\frac{3}{4}$ | $-\frac{1}{4}$ | $\frac{1}{4}$ | 0 | 0 | $-\frac{1}{4}$ | 1 | $-\frac{3}{4}$ | $\frac{3}{4}$ |
| $\frac{1}{2}$ | $\frac{1}{2}$ | 0 | 0 | $\frac{1}{4}$ | $-\frac{1}{4}$ | 0 | 1 | -1 | 1 |
| 1 | $\frac{3}{8}$ | $\frac{1}{8}$ | $-\frac{1}{8}$ | $\frac{3}{8}$ | $-\frac{3}{8}$ | $\frac{1}{8}$ | 1 | $-\frac{9}{8}$ | $\frac{9}{8}$ |
| ∞ | 0 | $\frac{1}{2}$ | $-\frac{1}{2}$ | $\frac{3}{4}$ | $-\frac{3}{4}$ | $\frac{1}{2}$ | 1 | $-\frac{3}{2}$ | $\frac{3}{2}$ |

Table 1: **Theoretical allometries.**
| c) Three-dimensional interactions ($\hat{\epsilon} = 5/6$) | | | | | | | | | |
| --- | --- | --- | --- | --- | --- | --- | --- | --- | --- |
| $\frac{r_{\beta\beta}}{r_{\alpha}}$ | $\hat{\alpha}$ | $\hat{\beta}$ | $\hat{t}$ | $\hat{p}$ | $\hat{R}$ | $\hat{r}$ | $\hat{h}$ | $\hat{n}$ | $\hat{w}$ |
| 0 | $\frac{5}{6}$ | $-\frac{1}{6}$ | $\frac{1}{6}$ | 0 | 0 | $-\frac{1}{6}$ | 1 | $-\frac{5}{6}$ | $\frac{5}{6}$ |
| $\frac{1}{4}$ | $\frac{2}{3}$ | 0 | 0 | $\frac{1}{6}$ | $-\frac{1}{6}$ | 0 | 1 | -1 | 1 |
| 1 | $\frac{5}{12}$ | $\frac{1}{4}$ | $-\frac{1}{4}$ | $\frac{5}{12}$ | $-\frac{5}{12}$ | $\frac{1}{4}$ | 1 | $-\frac{5}{4}$ | $\frac{5}{4}$ |
| ∞ | 0 | $\frac{2}{3}$ | $-\frac{2}{3}$ | $\frac{5}{6}$ | $-\frac{5}{6}$ | $\frac{2}{3}$ | 1 | $-\frac{5}{3}$ | $\frac{5}{3}$ |

**Table 2:**
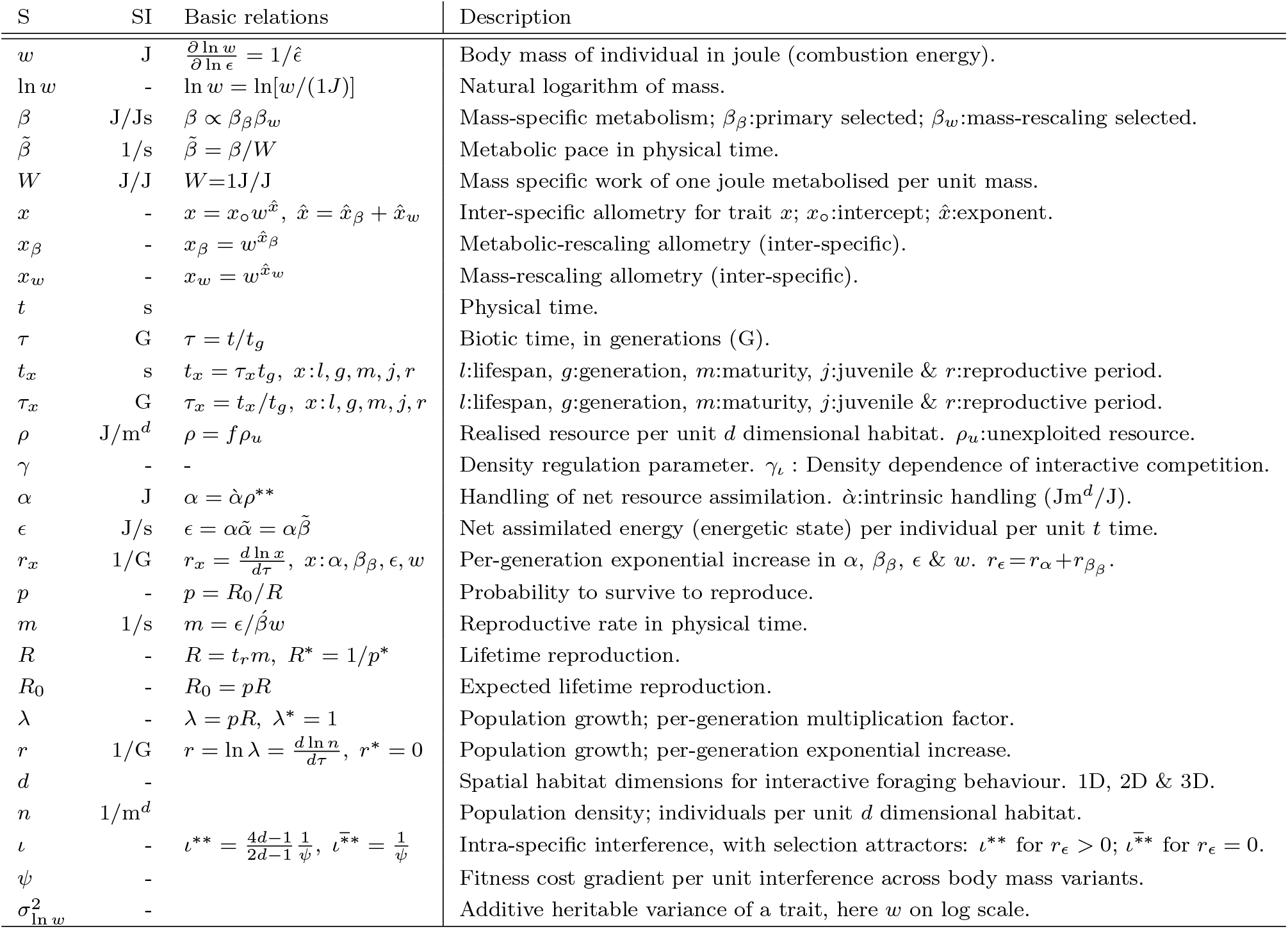
**Important symbols (S)** with SI units and basic relationships.

### 3.3 Mass in physical time

With equations for the evolution of handling, mass-specific metabolism and mass on a per-generation time-scale (eqns 29 and 31), and the associated allometric scaling with mass (Table 1), we have a predictive theory for life history evolution on the per-generation time-scale of natural selection. This evolution depends on the per-generation rate of change in mass (*r_w_* from eqn 6), with the remaining life history being a function of the evolving mass (Table 1), with a functional relationship that follows from the *r_β_β__/r_α_*-ratio and *d*.

To transform the predicted body mass trajectories to the physical time-scale of the fossil record, we use eqn 3 to scale the rate of change in mass with the evolutionary changes in generation time, with 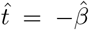. This predicts a natural selection time that dilates to the 1/4 power in 2D and the 1/6 power in 3D when there is no evolutionary increase in the mass-rescaling intercept for mass-specific metabolism. Natural selection time is body mass invariant when *r_β_β__/r_α_* = 1/2(*d* − 1), i.e., 1/2 in 2D and 1/4 in 3D, and it contracts to the −1/8 power in 2D and the −1/4 power in 3D for symmetrical unconstrained evolution where *r_β_β__/r_α_* = 1. At the *r_β_β__/r_α_* ≈ ∞ limit, where mass is increasing exclusively because of increased metabolism, the contraction of natural selection time occurs to the −1/2 power in 2D and the −2/3 power in 3D.

By scaling eqn 3 with the predicted 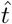 we obtain the 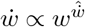 allometry, with the following exponent

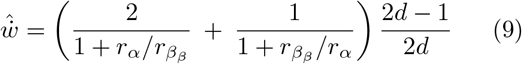

for the rate of change in mass in physical time (Table 1).

All of the predicted mass trajectories are log-linear on the per-generation time-scale of natural selection. Dependent on the *r_β_β__/r_α_*-ratio and *d*, the corresponding trajectories in physical time are bent downward 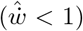 by a dilation of natural selection time, or upward 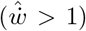 by a contraction. This is illustrated in Fig. 3, with the evolutionary trajectory being log-linear in physical time 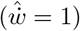 only when the *r_β_β__/r_α_*-ratio is 1/2(*d* − 1) (yellow trajectory).

**Figure 3:**
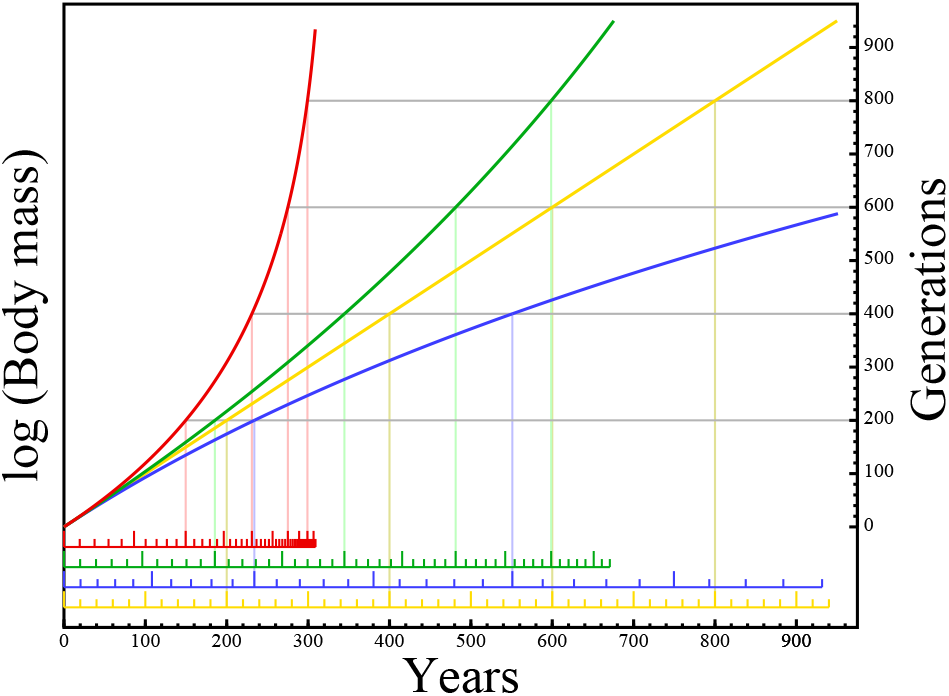
**Evolutionary bending** by contraction and dilation of natural selection time. A theoretical illustration of body mass evolution when the rate of exponential increase 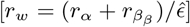 is constant on the per-generation time-scale of natural selection (2D interactions). The initial generation time is a year for all lineages, and the per-generation time-scale (right y-axis) is shown in physical time by the coloured x-axis. The evolutionary trajectories are the same for all lineages in generations, with the degree of bending in physical time following from the *r_β_β__/r_α_*-ratio. There is only a downward bend from time dilation when *r_β_β__/r_α_* = 0 (**blue**). Time dilation and contraction are equally strong (with no overall bend) when *r_β_β__/r_α_* = 1/2(*d* − 1) (**yellow**), while contraction with upward bending is dominating for unconstrained symmetrical evolution (*r_β_β__/r_α_* = 1, **green**) and extreme for within niche evolution with resource handling at an evolutionary optimum (*r_β_β__/r_α_* ≈ ∞, **red**).

These results are illustrated for placental mammals in Fig. 4. It shows the span of potential exponential trajectories from an estimated 125 gram ancestor at 65 ma (O’Leary et al. 2013) to a 10 tonnes terrestrial (2D) and 100 a tonnes pelagic (3D) mammal today. Included are also the 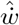 exponents for the evolutionary increase in mass (eqn 9).

**Figure 4:**
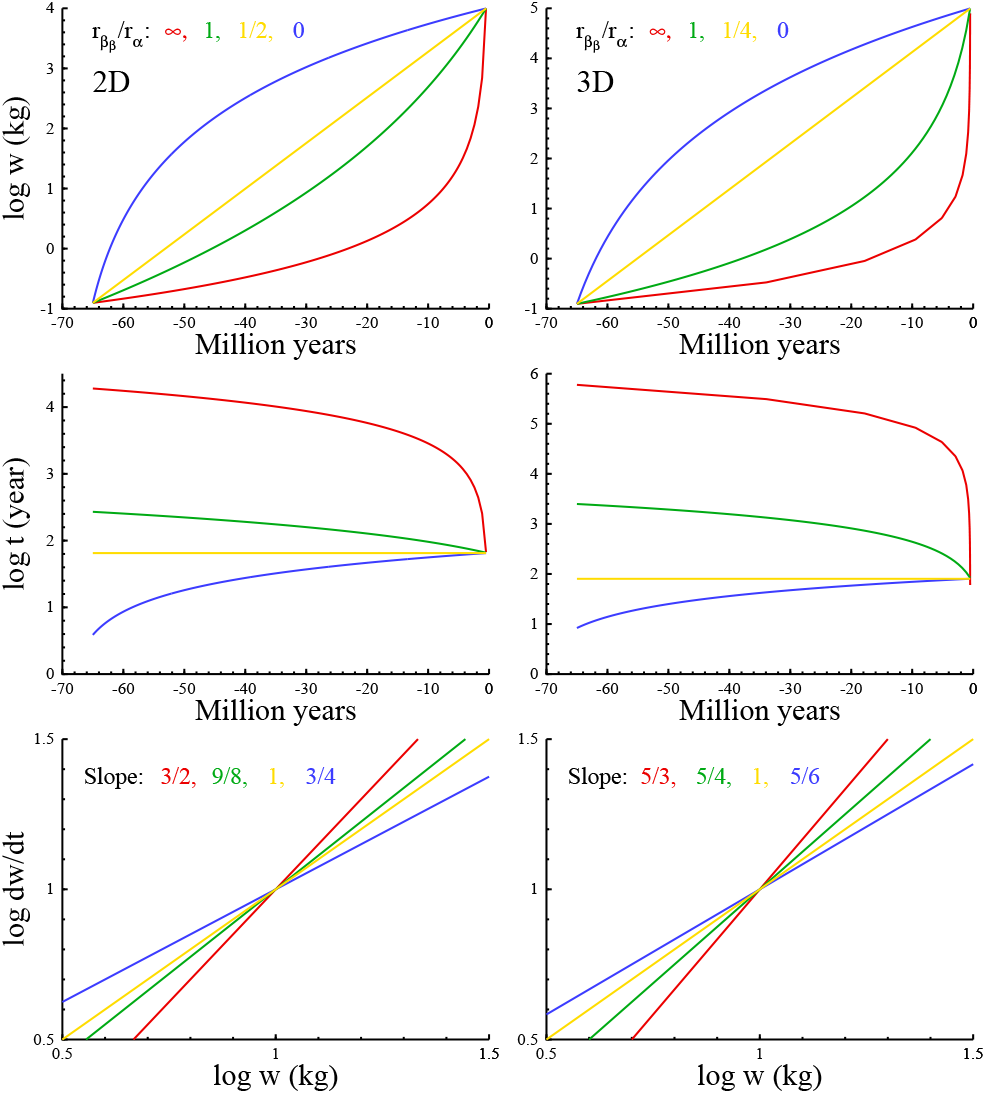
**Theoretical examples** for the evolution of maximum mammalian mass (w; **top**) and associated lifespan (*t*; **middle**) in physical time, given 2D and 3D interactions and constant rates of evolution on the per-generation time-scale. The *r_β_β__/r_α_*-ratio is estimated by the slope (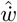 exponent, eqn 9) of the allometry 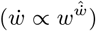 for the rate of evolutionary increase in body mass (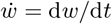, kg per million year) in physical time (**bottom**).

The shape of the trajectories depends on the *r_β_β__/r_α_*-ratio. When the ratio is zero it follows that net energy and body mass are increasing exclusively because of improved resource handling. This implies that there is no evolutionary change in mass-specific metabolism except for the decline that follows from the rescaling of the life history in response to the evolutionary increase in mass. The log trajectory is then levelling off over time with a 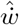 exponent of 3/4 or 5/6, dependent upon dimensionality. At the other extreme, the *r_β_β__/r_α_*-ratio is infinite, the energetic increase is exclusively due to enhanced metabolism, and the log trajectory is strongly convex with a 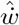 exponent of 3/2 or 5/3. In between, the trajectory is less convex when the *r_β_β__/r_α_* ratio is one, and it is linear when the ratio is 1/2 in 2D and 1/4 in 3D.

The predicted selections in Figs. 3 and 4 are compared with data in Figs. 5 and 6. The overall pairwise comparison between nine predicted and observed bending exponents 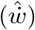 has a correlation coefficient of 0.98, with the evidence for each of the four types of selection being described in the subsections below.

**Figure 5:**
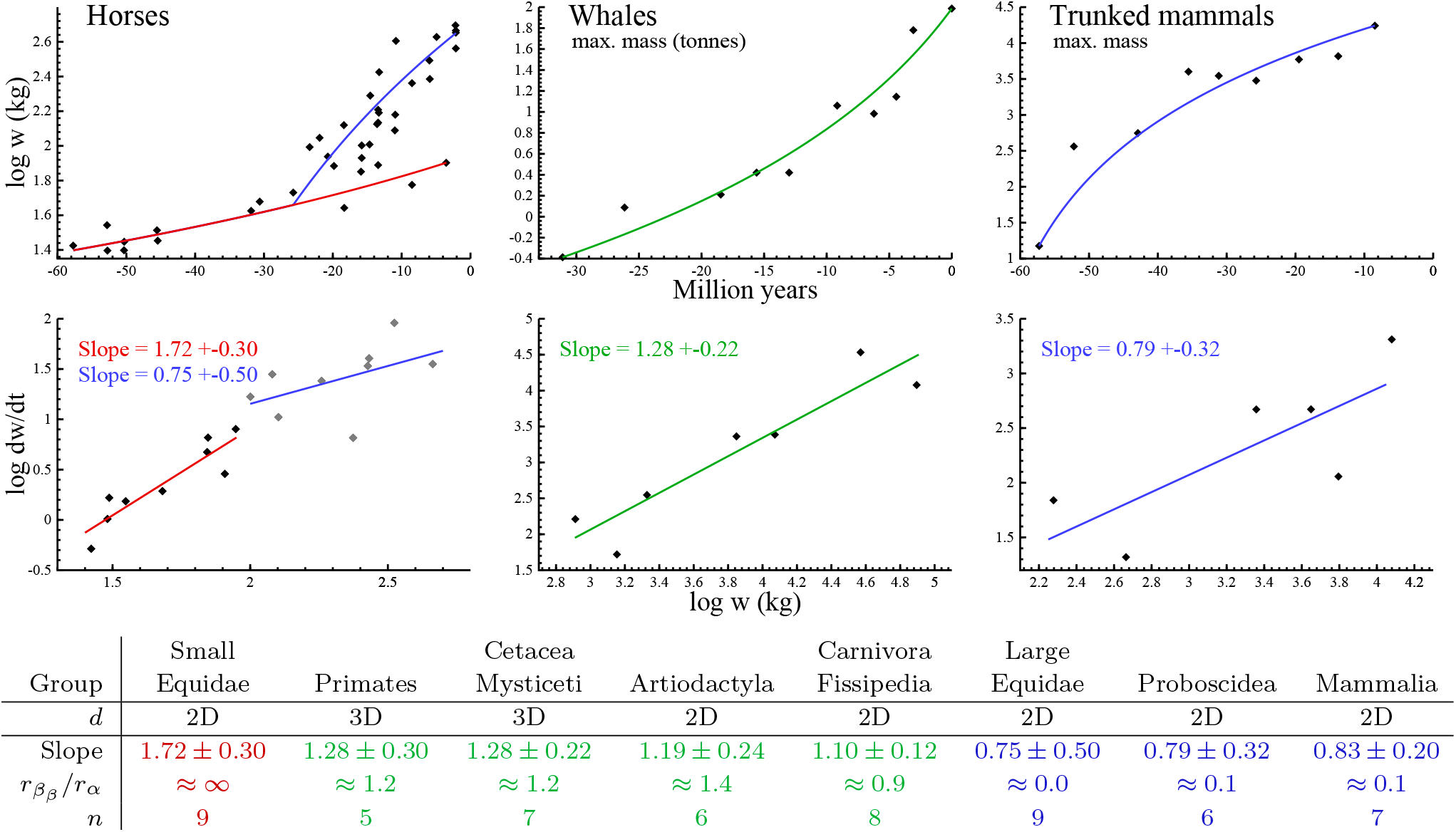
Mammalian evolution. Results for fossil horses (Equidae) and the maximum mass of other mammalian clades. **Plots:** Examples of theoretical (curves) and empirical (dots) body mass (w) trajectories (**top**). Curves are simulated from *r_β_β__/r_α_*-ratios that are calculated (by eqn 9) from the slope (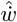-exponent and SE estimated by linear regression) of the allometry for the rate of increase in mass (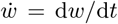, kg per million year) in physical time (**bottom plots**). **Table:** The estimated *r_β_β__/r_α_*-ratio approaches infinity for small fossil horses, indicating masses that are selected from an evolution increase in mass-specific metabolism (**red**). Most ratios for maximum mass are around one, with a similar increase in mass-specific metabolism and resource handling (**green**). With a ratio around zero, the increase in large horses and the maximum mass for trunked (Proboscidea) and terrestrial mammals is dominated by increased resource handling/availability (**blue**). No taxa showed log-linear evolution along a metabolic bound. 2D-3D classification from Witting (2017a). *n*: Number of data points in regressions, with data from MacFadden (1986), Smith et al. (2010) and Okie et al. (2013), see Supplementary Tables S1 to S4.

**Figure 6:**
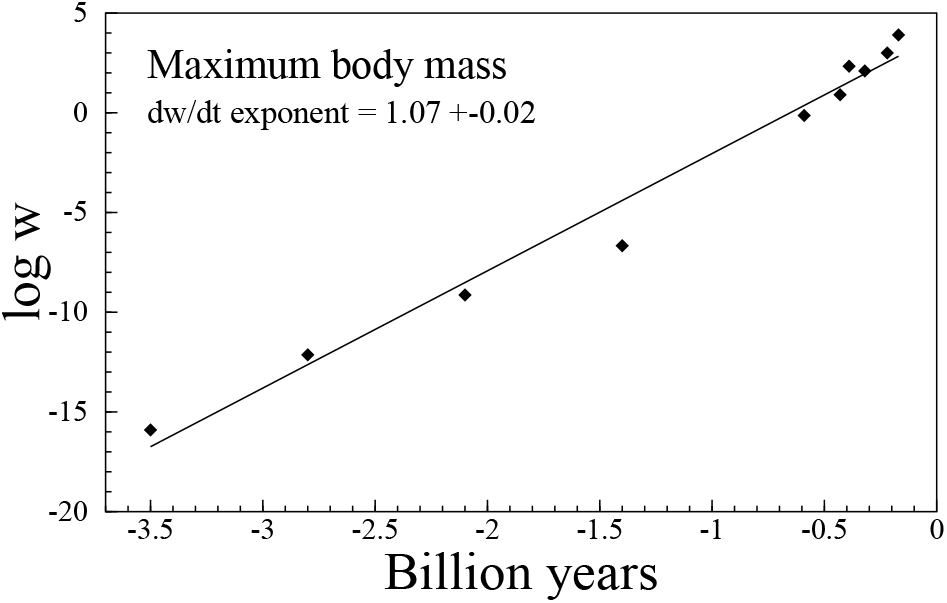
The maximum mass (length raised to third power) of mobile organisms over 3.5 billion years of evolution. The d*w*/d*t* exponent 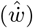 with SE is estimated by linear regression on the allometry for the rate of increase in mass in physical time. Data from Bonner (1965), Supplementary Table S5.

### 3.4 Evolution across niches 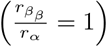

A symmetrical unconstrained selection across niches may be expected as a base case type of evolution for the maximum mass of taxonomic clades over time (Section 2.1). This evolution should generate an average *r_β_β__/r_α_*-ratio around one, with an associated 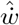 exponent of (6*d* − 3)/4*d*, i.e., 9/8 for 2D and 5/4 for 3D. The resulting trajectories are convex in physical time due to a time contraction where natural selection time evolves as 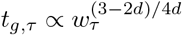 (green curves in Fig. 3 and 4).

This evolution is approximated for maximum mass in four out of five mammalian clades (Fig. 5, table). 2D 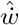 exponents around 1.13 are observed for 64 million years of evolution in terrestrial carnivores (Carnivora/Fissipedia), and 50 million years in even-toed ungulates (Artiodactyla). 3D 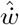 exponents around 1.23 are observed for 30 million years of evolution in whales (Cetacea & Mysticeti) and 55 million years in primates (Primates).

### 3.5 Fast evolution 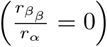

An average *r_β_β__/r_α_*-ratio around zero is expected for selection across niches when evolution in resource handling outruns evolution in mass-specific metabolism (Section 2.1). This is usually associated with fast body mass evolution, making it a candidate for the evolution of the largest species during evolutionary radiations. It has a 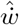 exponent of (2*d* − 1)/2*d*, and a concave trajectory in physical time due to a time dilation where the time-scale of natural selection evolves as 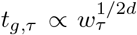 (blue curves in Fig. 3 and 4).

This evolution is not observed in whales, but the 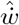 exponent is only slightly larger than the predicted 3/4 for the 2D-evolution of the maximum mass of trunked mammals (Probocidae) over 49 million years, and for the maximum mass of terrestrial mammals over 100 million years of evolution (Fig. 5). With a 2D 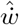 exponent of 0.75, it is indicated also for large horses during their evolutionary radiation over the last 25 million years.

### 3.6 Evolution within niches 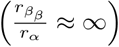

An average *r_β_β__/r_α_*-ratio that approaches infinity is expected for lineages that evolve within an ecological niche (Section 2.1). When resource handling adapts to the optimum of the niche and *r_β_β__/r_α_* → 0, it follows that net energy and mass can increase only by a selected increase in mass-specific metabolism, with *r_β_β__/r_α_* → ∞. Such lineages will have a 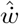 exponent of (2*d* − 1)/*d*, i.e., 3/2 for 2D and 5/3 for 3D. Their mass trajectories are strongly convex in physical time due to a time contraction where the time-scale of natural selection evolves as 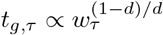 (red curves in Fig. 3 and 4).

While the larger horses during their evolutionary radiation after 25 ma show typical across-niche selection, we find that the evolution of smaller horses is driven by within-niche selection, with a 2D 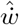 exponent of 1.72 ± 0.30 and a *r_β_β__/r_α_*-ratio around infinity (Fig. 5).

### 3.7 Evolution at metabolic limit 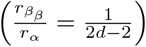

When mass-specific metabolism is selected along an upper bound we have that 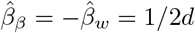, because the upward bend from the time contraction of metabolic acceleration is balanced against the downward bend from the time dilation of mass-rescaling. The result is an invariant mass-specific metabolism over time 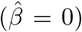, and a log-linear trajectory where 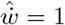 (yellow curves in Fig. 3 and 4). This evolution has a *r_β_β__/r_α_*-ratio of 1/2(*d* − 1), i.e., 1/2 for 2D and 1/4 for 3D.

The increase in maximum mass across the major life history transitions of non-sessile organisms over 3.5 billion years of evolution was found to have a 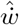 exponent of 1.07 ± 0.02 (Fig. 6) given data from Bonner (1965), and 1.07 ± 0.06 across selected maximum masses (Supplementary Table S6) from Payne et al. (2009). This apparent log-linear evolution on the largest macroevolutionary scale is supported by a body mass invariant mass-specific metabolism across major lifeforms from prokaryotes to mammals (Makarieva et al. 2005, 2008; Kiørboe and Hirst 2014; Witting 2017b). The latter invariance is observed not only across species distributions in general, but also for the maximum observed mass-specific metabolism across prokaryotes, unicellular eukaryotes, aquatic multicellular ectotherms, terrestrial ectotherms, and terrestrial endotherms (linear regressions on double logarithmic scale estimate slopes between −0.02 ± 0.01 and 0.01 ± 0.02, Supplementary Table S7).

## 4 Discussion

Fig. 6 plots the maximum limit on body mass evolution from the origin of the first self-replicating cells to whales; a scale that is covered by the life history theory of Malthusian relativity. Starting from inert replicators with no metabolism, the theory predicts not only the exponent of the limit trajectory, but also the evolutionary succession of lifeforms from a directional change in the primary selection of metabolism and mass with the selection increase in mass (Witting 2017a,b). This includes sexually reproducing multicellular animals as the fourth major lifeform in the succession. They are selected from larger unicells that are selected from small prokaryote-like self-replicating cells that are selected from inert replicators with no metabolism, no cell, and practically no mass.

The predicted major lifeforms follow as a monotonic function of the net assimilated energy that is available to an average individual in the population (Witting 2017a). With the current paper, I focused on the selection of mass and allometries from this energy, as it follows from the primary selection of resource handling and mass-specific metabolism. By comparing the predicted body mass trajectories with fossil data, I aimed for an independent confirmation of the underlying selection.

With mass-specific metabolism being selected as the rate of the handling that converts resource energy into replication (Witting 2017a,b), we predict an exponential increase in net energy and mass-specific metabolism on the per-generation time-scale of natural selection. I found this increase to explain not only the population dynamic feedback selection for an exponential increase in mass, but also allometric exponents for body mass trajectories in the fossil record.

### 4.1 A lack of fossil data

A direct test would compare the predicted exponents for body mass evolution in time with allometric exponents from fossil data. Yet these data are almost completely absent, as it is nearly impossible to estimate other life history traits than size from fossils.

There have been some attempts to estimate the age composition and maximal lifespan of fossil horses. Van Valen (1964), Hulbert (1984), and O’Sullivan (2005) used dental wear for age estimation. Yet, wear is rate dependent and there seems to be no straightforward way to convert dental wear rates in fossils into absolute age estimates. Other studies that use distinct dental wear-classes (Hulbert 1982) may be more promising, however, wear-classes may not necessarily represent year-classes (Hulbert 1984). For four species of fossil horses ranging back 18 million years (Hulbert 1982), the estimated lifespans varied from 70% to 75% of the expected given Kleiber scaling (as predicted by a 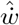 exponent of 0.75 in Fig. 5) and a 24 year lifespan for Burchell’s zebra (*Equus bruchelli*).

### 4.2 The curvature of evolution

With life history estimates for fossil species being too few for reliable estimates of allometries, I focused on body mass. The dependence of the allometric exponents on the *r_β_β__/r_α_*-ratio (Table 1) was extended to include the bending exponent 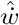. This allowed me to calculate the *r_β_β__/r_α_*-ratio and the allometric exponents from the curvature of body mass evolution, as estimated from fossils by the bending exponent for the rate of change in mass in physical time 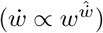.

Dependent upon the *r_β_β__/r_α_*-ratio, the bending exponent was predicted to change from 3/4 to 3/2 given 2D interactions, and from 5/6 to 5/3 given 3D. The limit trajectory across major lifeforms (Fig. 6) has a *r_β_β__/r_α_*-ratio of 1/2(*d* − 1) and a log-linear trajectory in physical time 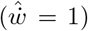. For the alternative case of within-niche selection with optimal resource handling, the predicted 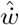 exponent is 3/2 in 2D, as approximated for small horses over 54 million years of evolution. This corresponds with an infinite *r_β_β__/r_α_*-ratio, and a mass that increases from a selection increase in mass-specific metabolism.

Symmetrical selection across ecological niches correspond with a *r_β_β__/r_α_*-ratio of one, relating to a body mass evolution where resource handling and mass-specific metabolism are equally important for the selected increase in mass. It is associated with 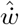 exponents of 9/8 in 2D and 5/4 in 3D, in agreement with the evolution of maximum mass in four out of five mammalian clades. A last option includes fast body mass evolution with a *r_β_β__/r_α_*-ratio of zero and 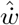 exponents of 3/4 in 2D and 5/6 in 3D. This is in agreement with the evolution of large horses, and maximum mass in trunked and terrestrial mammals.

It is important to note that the empirical exponents are somewhat uncertain as they follow from relatively few data points (table in Fig. 5). It is, e.g., somewhat surprising that the large size increase in whales is estimated with a symmetrical selection (*r_β_β__/r_α_* ≈ 1), instead of a fast selection that is dominated by an increase in resource handling or density (*r_β_β__/r_α_* ≈ 0).

### 4.3 Species distributions

The fossil trajectories in this paper is only a tiny fraction of the historical trajectories of evolution. While unlikely unique, the metabolic driven increase in small horses requires optimal resource handling. Moreover, the trajectories of maximum mass evolution represent only the upper boundaries of expanding species distributions. More generally, we expect not only time variation, but also lineages that have restricted and variable resource access due to inter-specific interactive competition. This generates not only evolutionary lineages with variable rates of increase, but also lineages with more stable masses, as well as lineages that decline in mass (Witting 2018).

Based on the selection framework of Malthusian relativity, Witting (2018) simulated the evolution of species distributions and inter-specific allometries across placental and marsupial terrestrial mammals. Starting from a single ancestor for each clade at the Cretaceous-Palaeogene boundary 65 ma, he simulated the evolution of present species distributions. The selection of different lineages in different niches illustrated how the life history of a single ancestor evolved into a species distribution with typical inter-specific allometries. Initial niche differentiations selected for a fast differentiation in net energy and mass, with a mass-rescaling selected Kleiber scaling where total metabolism increased to the 3/4 power of mass across species.

It was estimated that the selection of the major body mass variation from niche differentiation ceased around 50 to 30 ma, but also that the species distributions of body masses continued to evolve by an underlying background selection in mass-specific metabolism. This selection was strongest in placentals, where it bent the metabolic allometry over time and explained (Witting 2018) an observed curvature in the inter-specific allometry (Hayssen and Lacy 1985; Dodds et al. 2001; Packard and Birchard 2008; Kolokotrones et al. 2010; MacKay 2011). This created an approximate 3/4 exponent for the upper half of the size distribution, and an approximate 2/3 exponent for the lower half, providing a natural selection explanation for the 2/3 versus 3/4 controversy that has dominated the field of allometries for decades (e.g. Rubner 1883; Kleiber 1932; Heuser 1982; Feldman and McMahon 1983; Calder 1984; Dodds et al. 2001; White and Seymour 2003; Savage et al. 2004; Glazier 2010).

The feedback selection of Malthusian relativity is hence explaining both the evolution of individual trajectories and the evolution of species distributions; including Cope’s trend for a general increase in mass, the underlying body mass allometries, and their curvature in time and inter-specific space. This selection is somewhat in contrast to the more traditional view where an increase or decrease in mass is seen as being about equally likely, with both options having a multitude of potential causes (e.g., Schoener 1969; Stanley 1973; Gould 1988; Maurer et al. 1992; Bonner 2006; Brown and Sibly 2006; Clauset and Erwin 2008; Smith et al. 2010; Shoemaker and Clauset 2014). The selection of size has been argued to be so mechanistically diverse that it is best described by a neutral diffusion that produces an overall size increase by chance because evolution was initiated at a lower size limit (Stanley 1973; Gould 1988; McKinney 1990; Jablonski 1997; Clauset and Erwin 2008; Shoemaker and Clauset 2014).

Where a neutral diffusion may be sufficient for a statistical description of size distributions, it fails to explain rates of evolution in large versus small mammals (Baker et al. 2015), and the hypothesis is also insufficient from a mechanistic point of view. The quality-quantity trade-off [where parents can produce a few large or many small offspring from the same amount of energy (Smith and Fretwell 1974; Stearns 1992)] generates a constant frequency-independent selection for a continued decline in mass (Witting 1997, 2008, 2017a,b). This physiological selection for the near absence of mass implies, quite generally, that large animals can evolve only by a persistent frequency-dependent selection that is strong enough to outbalance the downward pull of the quality-quantity trade-off (Witting 2017b). The selection of species with non-negligible masses is a very active non-neutral process, where population dynamic feedbacks generate interactions between the frequency-independent selection of the physiology and the density-frequency-dependent selection of the intra-specific and inter-specific interactive competition.

### 4.4 Population dynamic feedback selection

It is intriguing that a natural selection that is continent and random at the basic level of genetic mutations, is found to predict existing lifeforms (Witting 2017b), the curvature of metabolic scaling in placental mammals (Witting 2018), and the curvature of body mass trajectories in the fossil record (this study). This is because population dynamic feedback selection evolves as a monotonic function from the very origin of inert replicating molecules. The feedback selection is a dissipative system that unfolds from the use of energy in self-replication, population growth, and intra-population interactive competition. With a steady influx of energy, there is a sustained evolution where a few general laws of selection are choosing a limited number of paths in the random space of mutations.

This implies a selection where a succession of life-forms, exponential evolutionary trajectories, and interspecific allometric variation follow more or less as a monotonic function of the selected variation in net energy across time and inter-specific space. A large fraction of the life history variation that were regarded as adaptations in the past are straightforwardly calculated from the selection balance between population growth and intra-population interactive competition.

### 4.5 Natural selection time

It is essential to recognise that selection operates on a per-generation time-scale, as observed for morphological (Lynch 1990; Gingerich 1993; Okie et al. 2013) and molecular traits (Martin and Palumbi 1993; Gillooly et al. 2005; Nabholz et al. 2008; Galtier et al. 2009; Broham 2011). With natural selection selecting mass-specific metabolism as the rate of biological processes, it follows that each species has its own time-scale of evolution that dilates and contracts relative to physical time dependent upon the evolutionary changes in metabolism and mass.

It is this dilation and contraction of the frequency of natural selection changes that bend the log-linear trajectories of exponential evolution in physical time. As the actual values of the predicted curvature agree with fossil data, the inclusion of evolutionary changes in natural selection time is essential for the interpretation of evolutionary changes in time.

## Acknowledgements

I want to thank those that collect fossils, estimate their body sizes, and publish the data. I am also grateful to the reviewers and editors that helped me improve the article.

## Appendix

### A Exponential increase

Earlier studies (Witting 1997, 2003, 2017a,b, 2018) have found large body masses to be selected by a population dynamic feedback selection, where the net energy that is allocated into reproduction generates population growth and interactive competition. The latter creates a positive correlation between net energy and mass because the larger than average individuals monopolise resources during interactive competition. The positive correlation is then selecting net energy into mass at the cost of reproduction and a continued increase in the density of the population.

As metabolism burns energy it follows that selection will optimise the physiological and ecological work that is essential for the life history. One implication of this (Witting 2017a), is a selection where the net energy

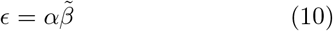

of the average individual (SI unit J/s) is selected as a product between the handling of resource assimilation [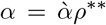; SI unit J, given by intrinsic handling 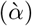 at the equilibrium resource density *ρ*\*\*], and the pace (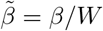; SI unit 1/s) of this process, with pace being selected as a proxy for mass-specific metabolism (*β*; SI unit J/Js), with *W* (SI unit J/J) being the mass-specific work of handling that is obtained by metabolising one joule per unit body mass, with mass (*w*; SI unit J) given as biotic (combustion) energy.

The average per-generation rate of replication

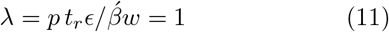

is one for populations that evolve over long periods of time, and it is proportional to net energy (*ϵ*), with *p* being the probability to survive to reproduce, 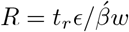 being lifetime reproduction (unitless number), *t_r_* (SI unit s) the reproductive period in physical time, and 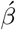 a unitless scaling that accounts for energy that is metabolised by the offspring (Witting 2017a).

Given the constraints of eqns 10 and 11, the per-generation selection gradients on the three resource assimilation parameters 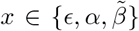 are unity on logarithmic scale

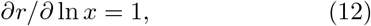

with *r* = ln λ. Then, from the secondary theorem of natural selection (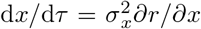, Robertson 1968; Taylor 1996) we expect an exponential increase

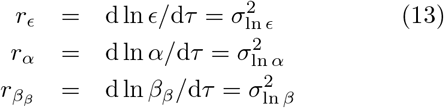

on the per-generation time-scale of natural selection, given unconstrained selection as defined by an invariant heritable variance 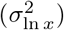. The rate of increase in net energy

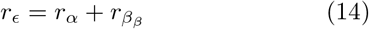

is the sum of the rates of increase in the two subcomponents of resource handling and metabolic pace. When selection is constrained, e.g., by an *α* that is approaching the selection optimum of a niche, or a *β* that is approaching an upper limit, it follows that the heritable variance will approach zero and that the rate of increase will cease.

The metabolic increase of eqn 13 is caused by the primary selection of metabolism (subscript *β*). This is to be distinguishes from the metabolic decline in the mass-rescaling component (*β_w_*) of mass-specific metabolism, which is a secondary selection response to the selection changes in mass (Witting 2017a).

Resource handling (*α*), on the other hand, is a pure primary selected parameter that generates net energy independently of the evolutionary changes in metabolism. It has a mass-rescaling exponent of unity (Witting 2017a) that reflects that an energetic increase in mass is caused exclusively by an increase in *α* for cases with no primary evolution in mass-specific metabolism. Although resource handling is defined to be *in principle* independent of the primary selection of metabolism, it does have an indirect metabolic-rescaling exponent 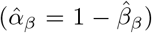 that reflects the importance of *α*, relative to *β*, for the generation of the net energy that is selected into mass (see eqn 25 in Witting 2017a).

Net energy in physical time (*ϵ*) is affected by mass-rescaling [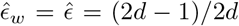, Witting 2017a]. Yet, net energy on the per-generation time-scale (which is the relevant time-scale for population dynamics and natural selection) is a pure primary selected parameter, with an allometric exponent of unity that reflects the conversion of net energy into mass. The mass-rescaling selection of mass maintains the current primary selected net energy by a dilation of natural selection time that balances the decline in net energy in physical time (Witting 2017a). The results is no direct, or indirect, metabolic-rescaling of net energy on either time-scale, with mass selected from net energy independently of the underlying cause (*α* versus *β*) for the generation of net energy.

The increase in net energy generates population growth with a density-dependent equilibrium, where eqn 83 in Witting (2017a) determines the level of interactive competition

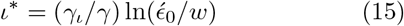

as a function of net energy, with *γ_ι_* being the density dependence of interference competition, *γ* the overall strength of density regulation, and 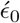 a measure of net energy on the per-generation time-scale. The population pressure of interference competition generates population dynamic feedback selection, where the resulting rate of increase in mass

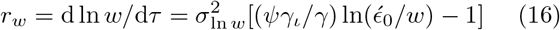

is a product between the selection gradient [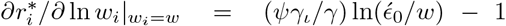, with subscript *i* denoting intra-population variation; from eqn 28 in Witting 2017b] and the heritable variance 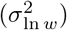, with *ψ* being the gradient in the fitness cost of interference per unit interference across the body mass variation in the population (see Witting, 2017a,b for details).

As the 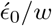 component of eqn 16 is invariant of mass (Witting 2017a), the expected increase is exponential, and it may be rewritten as

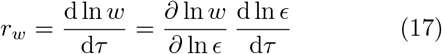

with an invariant selection relation

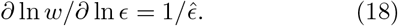

Hence, we have

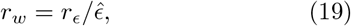

and a natural selection that defines mass

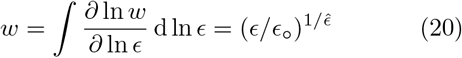

by the inverse of the net energy allometry

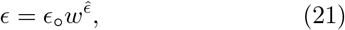

where 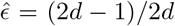 is the predicted exponent from Witting (2017a), *ϵ*_o_ the intercept, and *d* the spatial dimensionality of the intra-specific interactions.

By setting eqn 19 equal to eqn 16, and noting the constraint of eqn 11, we find the mass

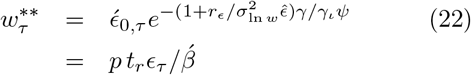

of the evolutionary steady state with an exponential increase in mass. Then, from eqn 16 we have

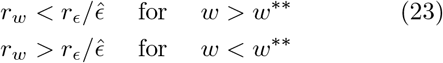

that confirm that the steady state is the unconstrained selection attractor for mass, with ** superscript denoting the attractor of unconstrained selection.

If we insert eqn 15 into eqn 16, and set 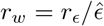 from eqn 19, we find the level of intra-specific interference competition

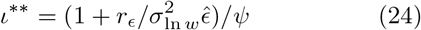

for an exponentially evolving mass to be somewhat higher than the level of interference for a stable mass at an equilibrium attractor 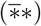 with an upper constraint on net energy, where 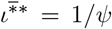 (from eqn 25 in Witting 2017b). As the parameters on the right hand side of eqn 24 are expected to be body mass invariant, the theoretically deduced allometries (Witting 1995, 2017a) apply for body mass evolution at steady state. As noted already, this implies an energetic exponent of 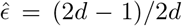, and with 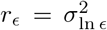 from eqn 13, we find that the level of interference reduces to

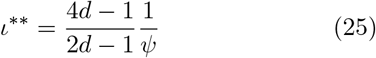

for the symmetrical case where 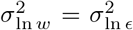. The position of this selection attractor of the steady state is shown on the selection integral in Fig. 1f in Witting (2017b).

Nearly all the allometries considered here and in Witting (2017a) describe variation in the average traits across evolved populations; either across species (Witting 2017a) or along an evolutionary lineage in time (current paper). Yet, the steady state defines also an important intra-specific allometry that describes the correlation between reproduction and mass across the individuals in the evolving population. To obtain this allometry, insert eqn 25 into the selection gradient 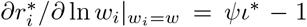 for the multicellular animal in Witting (2017b), and integrate over ln *w_i_* to find that the within population variation in the *pR*-product scale as

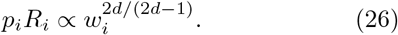

As this exponent is 4/3 and 6/5 for organisms with intra-specific interactions in two and three spatial dimensions, we find a reproductive rate that is about proportional to mass when survival is relatively invariant.

This proportionality is often observed in natural populations (Peters 1983), and it reflects a level of interactive competition that is so high that resource monopolization scales to the approximate second power of mass.

### B Allometries in time

The predicted exponential increase in metabolism and mass on the per-generation time-scale is an essential finding for evolutionary biology in itself. To understand the implication in detail, we need to transform the prediction into physical time to compare with fossil data. Hence, we need to predict the correlated evolution between mass and the per-generation time-scale of natural selection.

This correlation is given by the allometries in this subsection, with allometries being predicted not only for the generation time that is needed for the time-transformation of body mass evolution, but also for a larger set of traits that allow for more general predictions of life history evolution in time.

The exponents of the final allometries evolve as the sum of the partial correlations of the metabolic-rescaling and mass-rescaling that follow from the primary selection of metabolism and mass (Witting 2017a). To deduce this evolution we have from eqns 14 and 19 that

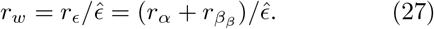

Now let

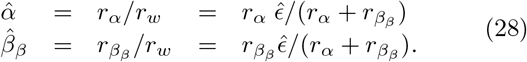

The exponential increase in *α, β_β, ϵ_* and *w* on the per-generation time-scale may then be expressed as a function of the exponential increase (*r_w_*) in mass, i.e.,

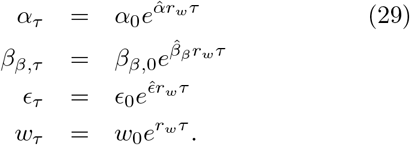

From these equations we can construct time allometries. For this, solve the body mass relation for time *τ* = ln(*w_τ_/w*_0_)/*r_w_*. Insert this time into the other relations and obtain

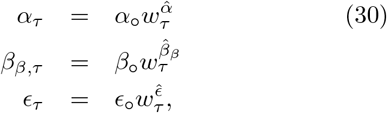

with intercepts *α*_0_ = *α*_0_/*w*_0_, *β*_o_ = *β*_*β*,o_/*w*_0_, and *ϵ*_o_ = *ϵ*_0_/*w*_0_. Hence, from eqn 28, 30, and 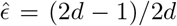, we find the following allometries for an evolutionary lineage in time

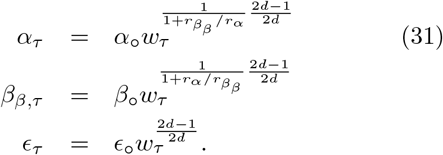

From the inverse relation between biotic time and metabolic pace, we find generation time to evolve with metabolic-rescaling as

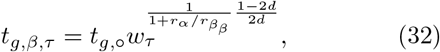

and mass-rescaling as

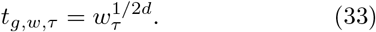

The mass-rescaling of eqn 33 dilates natural selection time as the body mass is increasing by natural selection. The mass-rescaling intercept, however, will decline by the metabolic-rescaling (eqn 32) that evolves by the primary selected mass-specific metabolism, and this causes a contraction of natural selection time. Whether natural selection time will actually contract or dilate depends on the level of metabolic-rescaling relative to mass-rescaling with the final allometry for generation time evolving as

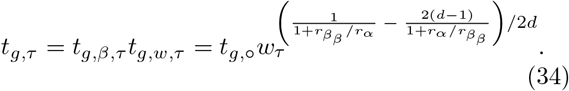

To transform the predicted trajectories to the physical time-scale of the fossil record, we have that the rate of change in mass in physical time is

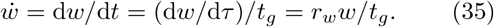

Then, from eqn 34 we obtain the following allometry

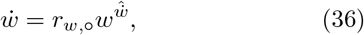

for the rate of evolutionary change in mass, where

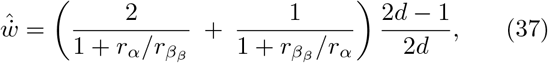

and *r*_*w*,o_ = *r_w_/t*_*g*,o_.

